# Amphibian mucus triggers a developmental transition in the frog-killing chytrid fungus

**DOI:** 10.1101/2022.01.21.477224

**Authors:** Kristyn A Robinson, Sarah M Prostak, Evan H Campbell Grant, Lillian K Fritz-Laylin

## Abstract

The frog-killing fungus *Batrachochytrium dendrobatidis* (*Bd*) grows within the mucus-coated skin of amphibians. Each lifecycle, *Bd* transitions from its motile, dispersal form to its sessile growth form in a complex process called encystation. Although encystation is critical to *Bd* growth, whether and how this developmental transition is triggered by external signals was previously unknown. We discovered that exposure to amphibian mucus triggers rapid and reproducible encystation within minutes. This response can be recapitulated with purified mucin, the bulk component of mucus, but not by similarly-viscous methylcellulose or simple sugars. Mucin-induced encystation does not require gene expression, but does require surface adhesion, calcium signaling, and modulation of the actin cytoskeleton. Mucus-induced encystation may represent a key mechanism for synchronizing *Bd* development with arrival at the host.

## MAIN TEXT

The parasitic chytrid fungus *Batrachochytrium dendrobatidis* (*Bd*) is decimating amphibian populations around the world (*1*–*4*). *Bd* has a biphasic life cycle, alternating between motile zoospores and sessile sporangia that produce new zoospores (*5*, *6*). Zoospores lack cell walls and swim rapidly through aquatic environments using a posterior flagellum and across solid surfaces using actin structures similar to those of human cells (*7*, *8*). *Bd* transitions from this motile dispersal form to its reproductive form by absorbing its flagellum, rearranging its actin cytoskeleton, and rapidly building a chitin-based cell wall—a process called “encystation” (*5*–*7*). After the resulting sporangium increases in volume by two or three orders of magnitude and undergoes rounds of mitosis without cytokinesis, it divides its cytoplasm into dozens of new zoospores. Daughter zoospores exit the sporangium through a discharge tube onto the amphibian skin and can then reinfect the same individual or find a new host (*5*).

*Bd* takes three to four days to complete its life cycle in pure culture, with zoospores taking up to 24 hours to encyst (*9*, *10*). How and whether this lifecycle changes in response to an amphibian host remains unknown. If *Bd* actively seeks out a host, it must do so prior to encystation as afterwards it can neither longer crawl nor swim (*6*, *7*). Moreover, if *Bd* retains its motility after finding a host, it risks leaving a productive growth site. This means that triggered encystation upon arrival at an amphibian host could increase colonization rates. Because *Bd* grows in amphibian skin that is coated in mucus (*5*, *11*)—a complex mixture of antimicrobial peptides, small molecules, and microbes that are embedded in a viscous matrix of heavily glycosylated proteins called mucins (*12*–*15*)—we hypothesized that amphibian skin mucus could trigger encystation.

To test our hypothesis that mucus induces *Bd* encystation, we imaged zoospores before, during, and for ten minutes following exposure to mucus isolated from wild green frogs (*Lithobates clamitans*). Because *Bd* zoospores rapidly swim out of the field of view, we adhered the zoospores to the imaging plane by coating the coverglass with the lectin concanavalin-A (ConA), a common surface coating for cell adhesion (*7*, *8*). Within minutes of exposure to amphibian mucus, adhered *Bd* zoospores retracted their flagella and rounded up—two clear signs of encystation (**Fig. 1A**, left and **Movie S1**). To quantify this encystation response, we counted the number of cells with retracted flagella over time and found that approximately 52% of mucus-treated cells encysted between two and seven minutes after exposure (mean time: 3.57 ± 0.19 minutes), while none of the control cells encysted during the same time period (**Fig. 1A, right**).

**Figure 1.**
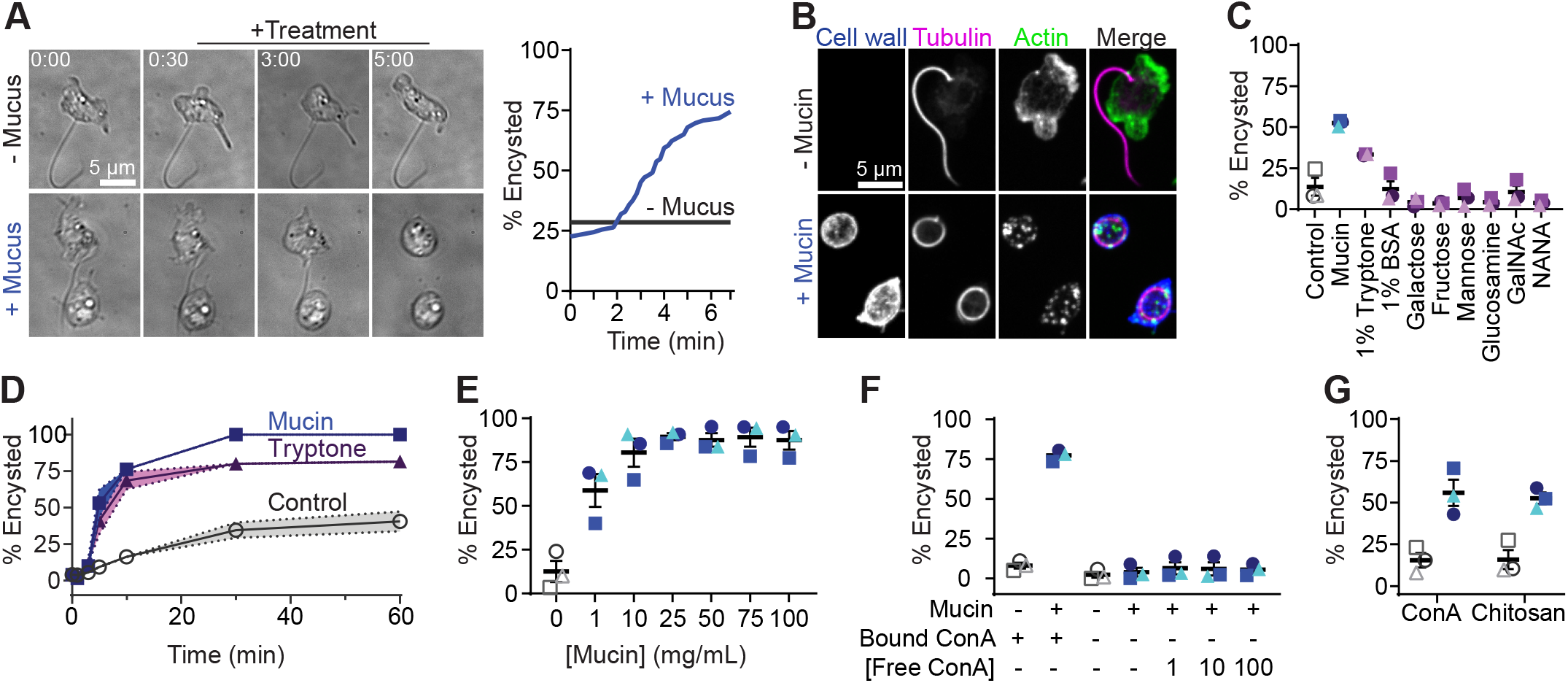
Adherent *Bd* zoospores encyst upon exposure to amphibian mucus or purified mucin. **(A)** Images of live *Bd* zoospores exposed to buffer (control) or amphibian mucus (min:sec). Right: Accumulation of encysted cells after mucus exposure. (**B**) Cells exposed to buffer (control) or 10 mg/mL purified mucin stained for cell wall (calcofluor white), flagella (tubulin tracker), and actin (phalloidin). (**C**) Percent cells encysted after five minutes of indicated treatment. (**D**) Time course showing percent cells encysted after treatment with buffer, 10 mg/mL mucin, or 1% tryptone. (**E**) Percent cells encysted after five minutes of treatment with the indicated concentration of mucin. (**F**) Percent cells encysted 5 minutes after addition of mucin or buffer (control) while adhered to a ConA coated surface (bound ConA) or in suspension with the indicated concentration of ConA in solution. (**G**) Percent cells encysted after five minute exposure to mucin or buffer (control) when adhered to ConA (left) or chitosan (right). (C-G) Open shapes represent control, filled shapes represent mucin (blue) or alternative treatment (purple). Means and standard errors of three independent biological replicates are shown. Significance (bold font for p<0.05) tested by ordinary one-way ANOVA (C,F) or unpaired t-tests (G).

We next tested if the encystation response could be recapitulated by exposure to mucins, a diverse family of heavily glycosylated proteins that form the viscous, bulk component of mucus (*13*–*15*). We added 10 mg/mL of commercially purified porcine stomach mucins to adhered cells, and saw a similar response (**Fig. 1B-D**). To confirm that the cells were encysting, we fixed zoospores five minutes after mucin exposure and stained them for cell wall material (calcofluor white), flagella (tubulin tracker), and actin (fluorescent phalloidin). While control cells lacked cell walls and had an actin cortex, pseudopods, and a posterior flagellum, mucin-exposed cells displayed all three hallmarks of encystation: actin patches, a retracted axoneme, and a cell wall (**Fig. 1B**). We quantified the percentage of cells that had each hallmark over time and found that, while only 50% of control cells encyst after one hour, nearly 100% of mucin-exposed cells are encysted within 30 minutes of mucin exposure (**Fig. 1D**). We saw equivalent results using mucin purified from either porcine stomach or bovine submaxillary gland (**Fig. S1B**).

The glycosyl chains of mucins are typically 2-12 monosaccharides long and include galactose, fructose, mannose, glucosamine (GlcN), N-acetylgalactosamine (GalNAc), N-acetylglucosamine (GlcNAC), and sialic acids, particularly N-acetyl-neuraminic acid (NANA) (*16*). We tested each of these constituents in our encystation assay, as well as a variety of other simple sugars, and found no response after five minutes (**Fig. 1C, Fig S1C**). To determine whether *Bd* may be responding to a physical property of hydrated mucin, we tested similarly-viscous methylcellulose (*17*), and saw no effect (**Fig. S1C**). We also tested altering the osmotic pressure of the extracellular environment by adding various concentrations of sorbitol, and again saw no effect (**Fig. S1C**). We next wondered if *Bd* is responding to the protein in mucin, and tested its response to another protein, bovine serum albumin (BSA), and again observed no response (**Fig. 1C**). Hypothesizing that *Bd* must be able to encyst in culture medium, we also tested the response to 1% tryptone—the media in which we grow *Bd*—and found that this treatment induced a partial encystation response (33.33% ± 0.2728, p=0.0034, **Fig. 1C**).

To measure the speed of the encystation response to mucin and tryptone, we conducted an encystation time course between one and 60 minutes after exposure to buffer (control), 10 mg/mL mucin, or 10 mg/mL tryptone (1%) (**Fig. 1D**). We found that while all (100% ± 0) of mucin treated cells encysted after 60 minutes, fewer tryptone-treated cells encysted (81.5% ± 4.1). The kinetics of the response also differed slightly; mucin-treated cells reached half maximum encystation by three minutes, while tryptone took five minutes. We next measured the dose-response of mucin- and tryptone-induced encystation at five minutes. We found that while the encystation response to both treatments were dose-dependent, the maximum response to tryptone resulted in the encystation of only 18.7% ± 3.38 of cells (**Fig. S1D**, 10 mg/mL tryptone), while mucin could induce encystation in 89.57% ± 1.96 (**Fig. 1E**, 25 mg/mL mucin). Given the speed and near completeness of mucin-induced encystation, we also tested whether the kinetics of mucin-induced encystation were also concentration-dependent, and found that treatment with 100 mg/mL mucin decreased the time to half maximum encystation to less than a minute (**Fig. S1E**). Because *Bd* encysted within 5 minutes of treatment with frog mucus, and previous estimates of mucin concentrations of 10-20 mg/mL in the skin mucus of aquatic animals (*18*), we chose a five minute exposure of 10 mg/mL mucin as our standard for the remaining experiments.

We next tested whether surface adhesion was necessary for mucin-induced encystation by exposing zoospores to mucin in the presence and absence of a ConA-coated surface, and found that only cells adhered to ConA-coated surfaces encysted in response to mucin (p=<0.0001, **Fig. 1F**), even when we added various concentrations of ConA in suspension. To determine whether mucin-induced encystation is specific to ConA-mediated adhesion, we first needed to identify other surfaces to which *Bd* could attach. By testing a panel of other possible surface coatings (**Fig. S1F**), we found that *Bd* adheres equally well to chitosan (a polymer derived from chitin (*19*)), and only slightly to polyethyleneimine (another commonly-used cell adhesive (*20*)) and mucin. We confirmed that *Bd* tightly binds ConA and chitosan using interference reflection microscopy to visualize the region of the cell in close contact with the coverslip (*21*) (**Fig. S1G**). *Bd’s* adhesion to ConA and chitosan may be an active process as heat-killed cells fail to adhere to either substrate (**Fig. S1F)**. We next compared the ability of ConA and chitosan to facilitate mucin-induced encystation and found that both surface treatments worked equivalently (p=0.0112 and p=0.0055, respectively, **Fig. 1G**). Together, these data indicate that mucus-induced encystation can be recapitulated by purified mucin, but not simple sugars, and requires surface adhesion.

Previous reports suggested that encystation of *Bd* may not require gene expression (*22*). To test whether mucin-induced encystation involves gene expression, we treated cells with 150 μg/mL of cycloheximide for 10 minutes prior to encystation. Although this concentration (533 μM) is sufficient to block downstream growth and development (**Fig. S2A**), cells were still able to robustly encyst upon exposure to mucin (p=0.0002, **Fig. 2A-B**), indicating that mucin-induced cell wall assembly requires no new protein synthesis. This is similar to encystation in other chytrid species (*23*, *24*).

**Figure 2.**
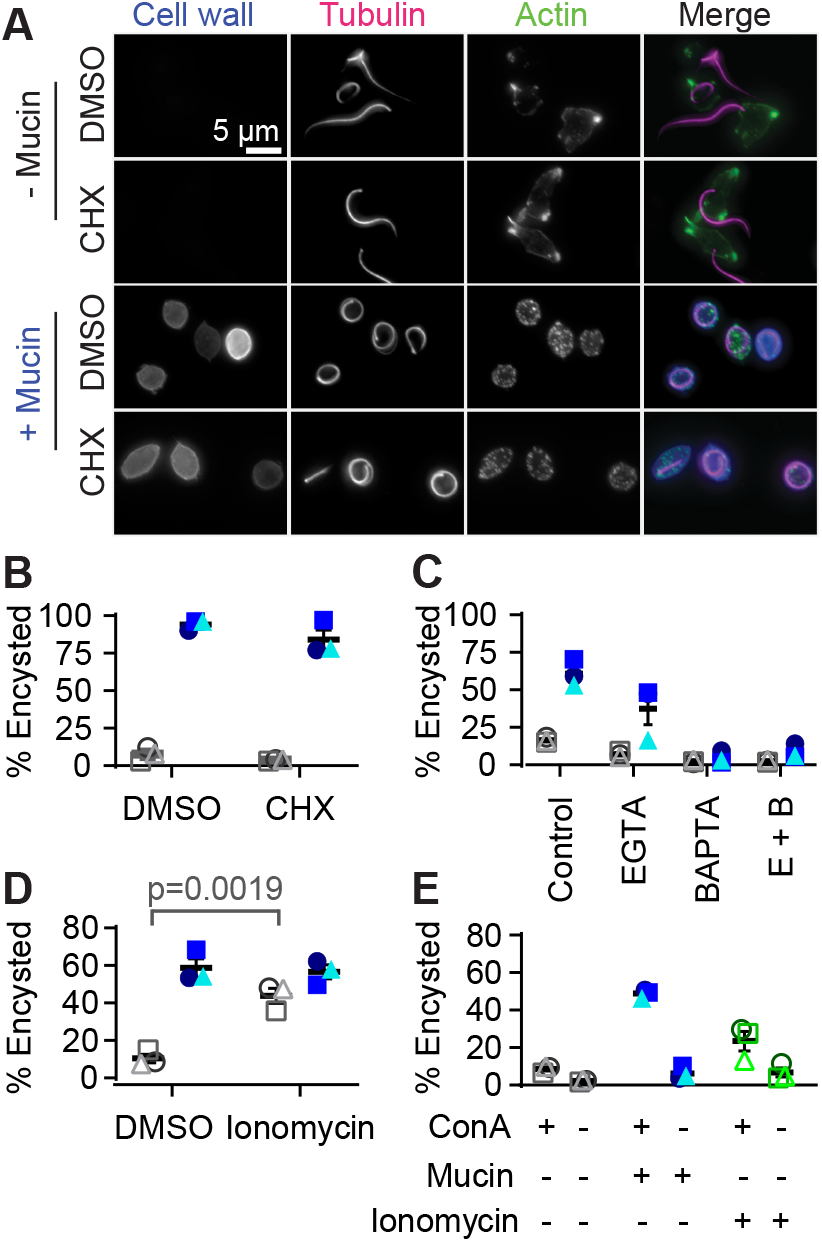
Encystation is driven by calcium signaling, not protein translation. (**A**) Cells exposed to buffer (control) or 10 mg/mL mucin and DMSO or cycloheximide (CHX) stained for cell wall (calcofluor white), flagella (tubulin tracker), and actin (phalloidin). (**B**) Percent cells encysted after DMSO (left) or cycloheximide (CHX; right) treatment upon exposure to control buffer or mucin. (**C-D**) Percent encysted cells treated with the indicated (C) calcium chelator, (D) ionophore, or (C-D) DMSO carrier control five minutes after adding mucin or control buffer while adhered to a ConA coated surface. (**E**) Percent cells encysted after addition of control buffer, mucin, or ionomycin while adhered to a ConA coated surface or in suspension. Filled shapes represent mucin (blue), open shapes represent the control (gray) and ionomycin (green), and each shape corresponds to data from three independent biological replicates. Significance (bold font or p<0.05) tested by unpaired t-test (B,D) or ordinary one-way ANOVA (C,E). Means and standard error of the mean of three biological replicates are shown.

Given its speed, we hypothesized that calcium signaling may be coordinating the events that comprise encystation. We therefore pre-treated ConA adhered zoospores with the calcium chelators EGTA and/or BAPTA for ten minutes prior to mucin-exposure and observed a 23.6 - 56.2% reduction in encystation (p=0.0002, p=<0.0001, and p=<0.0001 for the combination, **Fig. 2C, S2B**), suggesting that calcium flux is necessary for encystation. We next tested whether ionomycin—a calcium ionophore that transports calcium across membranes—can induce encystation. Indeed, treatment with 3 μM ionomycin induced encystation in the absence of mucin (p=0.0019, **Fig. 2D**). Further testing indicated that ionomycin does not subvert the need for adhesion (**Fig. 2E**), indicating that once cells are adhered to a surface, calcium flux is sufficient to induce encystation. Finally, we tested whether either mucin or ionomycin could induce encystation in the non-parasitic chytrid model system *Spizellomyces punctatus*, and found that they did not encyst in response to either (**Fig. S2C**). Together, these data indicate that mucin-induced encystation is a calcium-dependent response that may be limited to *Bd*.

Fungal cell wall assembly is largely driven by enzymes that are delivered in vesicles to the plasma membrane (*25*). Because zoospores have an actin cortex that may block vesicles from docking with the plasma membrane (*7*), we tested whether interfering with the actin cytoskeleton would modulate mucin-induced encystation. To this end, we treated adhered zoospores with a panel of small molecules that have previously been shown to modulate actin polymerization and turnover in *Bd* (*7*), as well as blebbistatin and Cytochalasin D that we here show alter *Bd* actin (**Fig. S3A-B**), along with their vehicle controls. Although we observed no statistically significant effect of any actin inhibitor on mucin-induced encystation (**Fig. 3A, Fig. S3A-C**), we found that latrunculin—a compound that sequesters actin monomers and results in complete loss of polymerized actin in *Bd* zoospores (*7*)—resulted in an 16.47 ± 2.77% increase in encystation in non-mucin treated cells (p=0.0040, **Fig. 3A**). Similarly, treatment with 10 μM jasplakinolide, a concentration that results in actin cortex reduction in *Dictyostelium* (*26*), also increased encystation in non-mucin treated cells (12.80% ± 3.28 vs 33.90% ± 0.81, p=0.0033). Together, these data indicate that actin depolymerization promotes encystation. This may be by way of depolymerizing the actin cortex to allow vesicle release, a model consistent with vesicular delivery in mammalian cells with an actin cortex similar to that of *Bd* (*7*, *27*).

**Figure 3.**
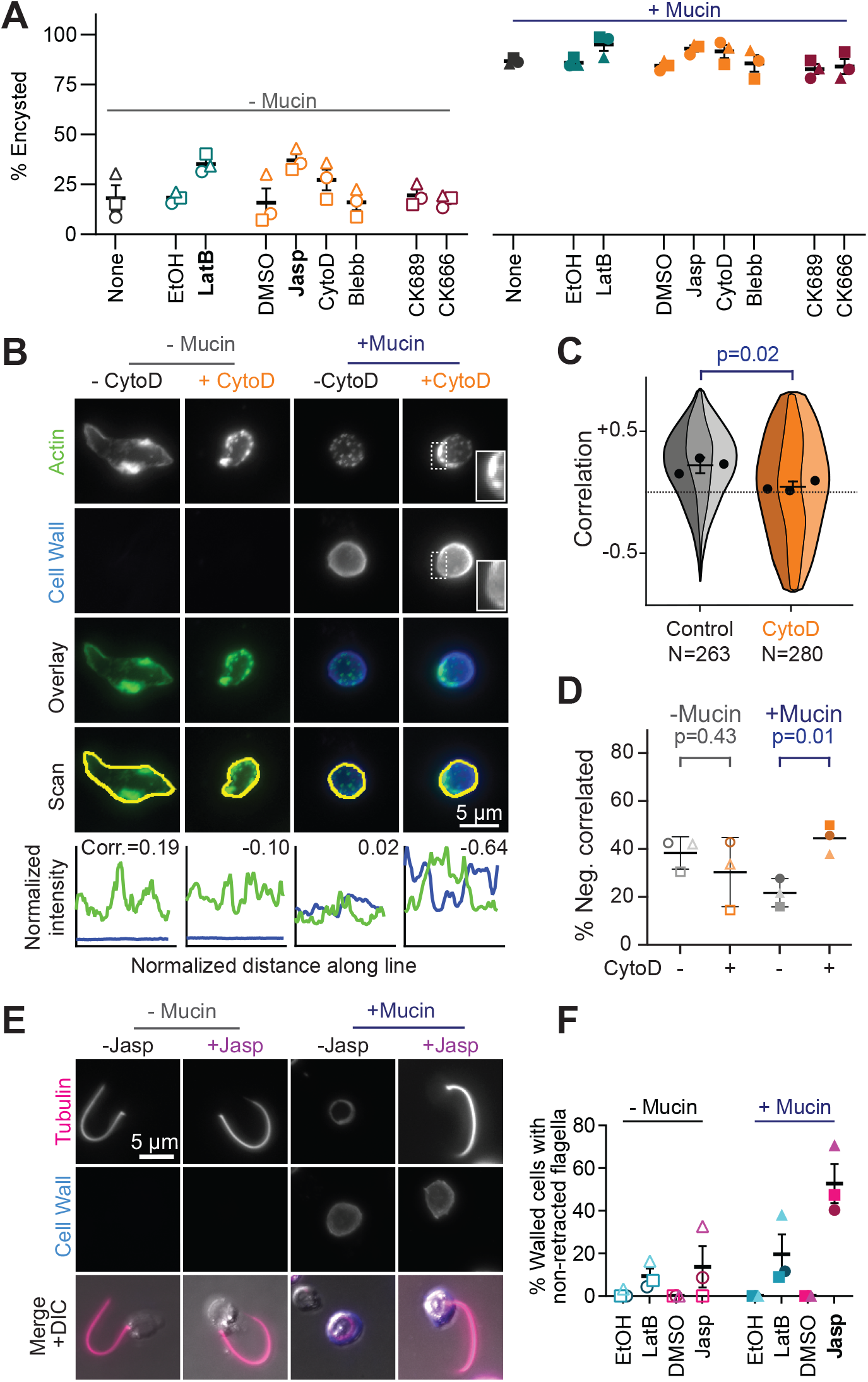
Actin cortex disassembly is associated with cell wall assembly. (**A**) Percent cells encysted when exposed to buffer (left) or mucin (right) after indicated treatment. (**B**) Representative cells treated with DMSO or Cytochalasin D (CytoD), exposed to buffer or mucin, and stained for actin (phalloidin) and cell wall (calcofluor white). Graphs show normalized fluorescence intensity of actin (green) and cell wall (blue) along yellow lines with associated Pearson’s correlation coefficient. (**C**) SuperViolin plot (*29*) of correlation coefficients of actin and cell wall intensity in mucin-exposed cells treated with DMSO (gray) or CytoD (orange). N = number of cells quantified by line scans; p-values calculated using unpaired t-test, shades indicate independent replicates. (**D**) Percent cells with negatively correlated actin and cell wall intensities upon exposure to buffer or mucin after treatment with DMSO or CytoD. (**E**) Representative cells treated with DMSO or jasplakinolide (Jasp), exposed to buffer (control) or mucin, and stained for cell wall (calcofluor white) and tubulin (tubulin tracker). (**F**) Percent of walled cells with non-retracted flagellum upon exposure to buffer or mucin after indicated treatment. LatB; Latrunculin B, Jasp; Jasplakinolide. (A, F) Bold font indicates p<0.05 by unpaired t-test of independent replicates (shapes).

To test this hypothesis, we analyzed mucin-encysted cells treated with Cytochalasin D, a inhibitor that binds to the growing ends of actin filaments and both stabilizes and prevents further elongation. Although not statistically significant, Cytochalasin D treatment resulted in higher numbers of encysted cells in both mucin-treated and control conditions (difference of 14.53% ± 8.99, p=0.1812 and 15.63% ± 6.82, p=0.0835 respectively, **Fig. 3A**). Interestingly, encysted cells treated with Cytochalasin D appeared to have retained cortical actin in some areas that also appeared reduced in cell wall content (**Fig. 3B**). To quantify this relationship, we measured actin and cell wall staining intensity along the periphery of the cell and calculated the Pearson’s Correlation Coefficient (*28*). Cytochalasin D treatment resulted in a reduction in the average correlation coefficient (**Fig. 3B-C**), as well as an increase in the percent of cells with a negative correlation between cortical actin and cell wall staining in mucin-treated cells (**Fig. 3D**). Taken together, these results suggest that reduction in cortical actin is associated with increased cell wall assembly.

During these experiments, we noted that jasplakinolide-treatment resulted in cell wall assembly in cells that retained non-retracted flagella (0% ± 0 vs 46.25% ± 10.12, p=0.0103, **Fig. 3E**). To determine if this effect is specific to jasplakinolide, we measured the number of cells with cell walls and non-retracted flagella in latrunculin-treated cells treated and found a similar effect (0% ± 0 vs 17.85% ± 8.02, p=0.0902, **Fig. 3F**), suggesting that perturbation of actin dynamics can uncouple flagellar retraction from cell wall assembly during mucin-induced encystation.

Together, our data indicate that exposing *Bd* zoospores to amphibian mucus or purified mucin triggers a rapid encystation response that does not require new gene expression, but does require surface adhesion. Moreover, we show that this mucin-induced encystation is controlled by calcium. We also show that actin cortex disassembly is associated with cell wall accumulation. These data are consistent with the following model (**Fig. 4**): (1) zoospores sense mucin and, (2) respond by inducing a calcium signaling cascade that results in the depolymerization of the actin cortex and (3) subsequent delivery of cell wall synthesis enzymes to the plasma membrane for (4) cell wall assembly. This model could explain how *Bd* zoospores transition from the motile dispersal form to their growth form at the appropriate time and place to increase the probability of successful host colonization.

**Figure 4.**
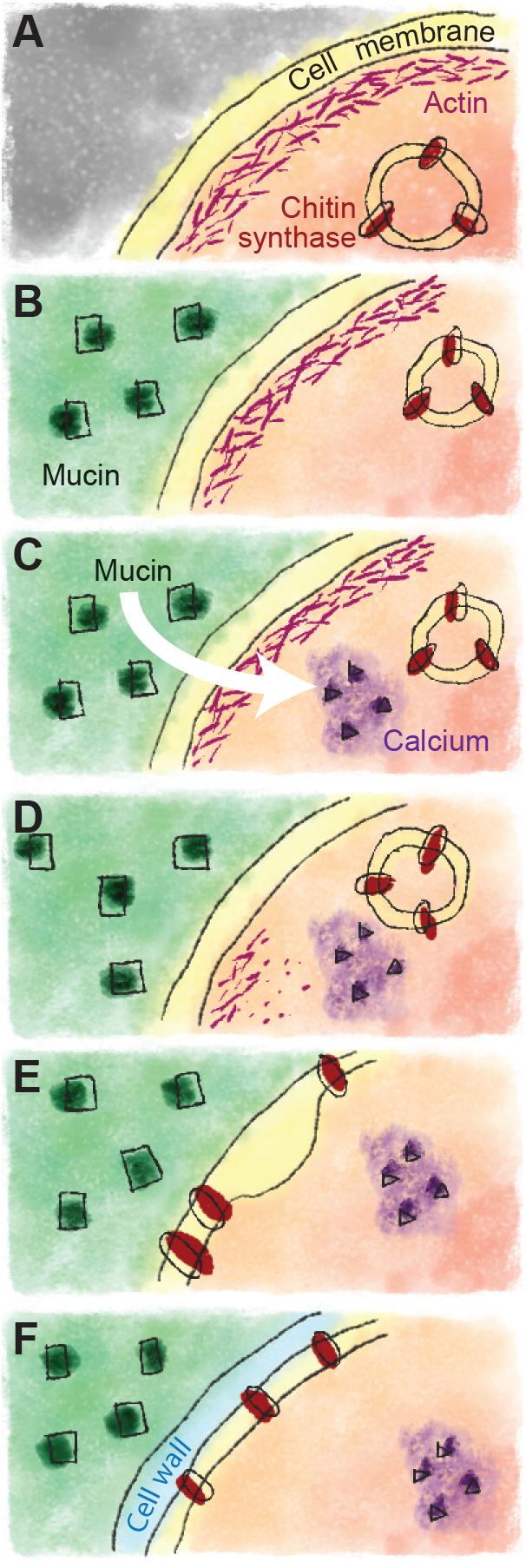
Model for mucin-induced encystation. We propose that: (**A**) *Bd* zoospores contain chitin synthases or other cell wall synthases in intracellular vesicles that are blocked from the plasma membrane by cortical actin. (**B**) Zoospores sense arrival at the host by the presence of mucin that (**C**) induces a calcium spike. (**D**) The subsequent depolymerization of cortical actin (**E**) allows chitin synthase-containing vesicles to dock with the membrane, which (**F**) results in the synthesis of cell wall material.

## MATERIALS AND METHODS

### Amphibian mucus collection

Northern green frog adults (*Lithobates clamitans*) were captured by dipnet, and transferred using new latex gloves to individual plastic bags. Individuals were measured in the bag and released, leaving behind mucus secretions on the inside surfaces of the bag. (Measurements were made for other, non-associated experiments.) Bags were transported from the field in a cooler on ice and stored at −20°C. Just before use, bags were thawed at room temperature, and mucus squeezed into the corner of each bag using a technique similar to that typically used to extrude the final remnants of toothpaste from a nearly-empty tube, and pipetted out of each corner of each bag and transferred into 1.5 mL tubes for immediate use.

### Cell culture

Age-matched zoospores of *Batrachochytrium dendrobatidis* strain JEL423 (*Bd*) were grown and collected as previously described (*30*). Briefly, semi-synchronized cultures were grown in 1% tryptone (w/v; Sigma, T7293) at 24°C in tissue culture-treated flasks (Fisher, 50-202-081). The adhered sporangia were then washed thrice with culture media, and incubated in fresh media for 2 hours 24°C. The zoospores that were released during this incubation were then collected by centrifugation at 2,500Xg. *Spizellomyces punctatus* (ATCC strain 48900, *Sp*) was grown on K1 Carb/Tet (1L; 0.6 g peptone, 0.4 g yeast extract, 1.2 g glucose, 15 g agar if plates; 50 μg/mL Carb/Tet) plates and zoospores collected by flooding each plate with 1.5 mL DS solution (50 mM KH_2_PO_4_, 50 mM K_2_HPO_4_, 50 mM (NH_4_)_2_HPO_4_, 50 mM MgCl_2_, 50 mM CaCl_2_). Zoospores were then transferred directly to a syringe and filtered using Whatman #1 (Fisher, 098051G).

### Quantification of encystation

8-well cover-glass bottom dishes (Eppendorf, 0030742036) were plasma cleaned and immediately coated with 0.5 mg/mL Concanavalin A (Sigma, C2010). Synchronized zoospores were washed three times with Bonner’s salts (10.27 mM NaCl, 10.06 mM KCl, 2.7 mM CaCl_2_ in MilliQ water), and added to individual wells. 100 μL volume of either Bonner’s Salts or amphibian mucus was then added to each well while imaging.

Mucin (Mucin from porcine stomach, type III, Sigma, M1778; Mucin from bovine submaxillary glands, Sigma, M3895) was resuspended in Bonner’s Salts for 2 hours at room temperature with shaking. Particulates were then removed by centrifugation at 15000 RCF for 2 minutes. Galactose (Sigma, G0750), fructose (Sigma, F0127), mannose (Sigma, M6020), N-acetylgalactosamine (Sigma, A2795), glucosamine (Sigma, G4875), and N-acetylneuraminic acid (Sigma, A0812) were dissolved in Bonner’s Salts at 20 mg/mL. Solutions of α-Lactose monohydrate (Sigma, L2643-500), sucrose (Fisher, S5-500), dextrose (Sigma, D9434-250g), starch (Alpha Aesar, A11961), N-acetylglucosamine (Sigma, A4106) and methyl cellulose (Sigma, M7140) were made in Bonner’s Salts at 50 mg/mL. 1M solutions of Glycerol (Sigma, G5516) and D-Sorbitol (Sigma, S1876) solutions were made in Bonner’s Salts. 1% BSA (Fisher, BP1600-1) was made the day of the experiment in Bonner’s Salts. Except when indicated, adhered synchronized zoospores in 100 μL Bonner’s salts were treated with 100 μL of each solution 5 minutes before fixation. Zoospores in suspension were treated with mucin supplemented ConA, 100 μg/mL, 10 μg/mL or 1 μg/mL.

### Adhesion Assays

96-well plates (MatriPlate, MGB096-1-2-LG-L) were plasma cleaned and coated with one of the following: Bonner’s Salts, Concanavalin A, 0.1% polyethyleneimine (w/v; Sigma, P3143), 0.1% poly-L lysine (w/v; Sigma, P8920), 100 μg/mL fibronectin (Sigma, F4759), 1:1000 keratin (Sigma, K0253), 0.05% chitosan (w/v; Sigma, 448869), and 10 mg/mL mucin. Wells were incubated for 1 hour and washed three times with Bonner’s. Adhered synchronized *Bd* zoospores were washed five times with 100 μL Bonner’s Salts using a 12-channel pipette. To heat kill, synchronized zoospores were placed in a 65 °C water bath for 1 minute and immediately placed on ice for 1 minute.

### Small molecule inhibitors

Adhered synchronized zoospores were treated with either Bonner’s Salts, DMSO or 150 μg/mL Cycloheximide (to inhibit protein translation; Santa Cruz, SC 3508-b), 1 μM Latrunculin B (to sequester actin monomers; Millipore, 4280201MG), or an equal volume of ethanol, 100 μM CK666 (to inhibit the Arp2/3 complex; Calbiochem/Sigma, 182515), 100 μM CK689 (an inactive control for CK666; Calbiochem/Sigma, 182517), 100 μM Cytochalasin D (to cap actin filaments; Gibco/Life Technologies, PHZ1063), 10 μM Jasplakinolide (to stabilize actin filaments; Molecular Probes/Invitrogen, J7473), 5 μM Blebbistatin (to inhibit myosin II; Cayman Chemical, 13891), or equal amounts of DMSO for 10 minutes. The cells were then treated with either Bonner’s Salts or mucin for 5 minutes before fixation. To test the long-term effect of Cycloheximide, synchronized zoospores were added to a 6-well TC plate (NEST, 703001). Wells were treated with concentrations of Cycloheximide suspended in 1% Tryptone for three days: 0 μg/mL, 50 μg/mL, and 150 μg/mL. On the third day, the wells were imaged using the 10X objective. To test the efficacy of phalloidin labeling on Jasplakinolide-treated cells, zoospores were treated with 10 μM Jasplakinolide before and after fixation, then labeled with phalloidin.

### Calcium signaling

Bonner’s Salts was used to make the solutions of 3 μM Ionomycin (Sigma, I0634) and DMSO carrier control. Two derivatives of Bonner’s salts solutions were also used: one which replaced calcium chloride with magnesium chloride (referred to as Mg in **Fig S2**) and without calcium or magnesium chloride (referred to as NDC, or No Divalent Cations, in **Fig S2**). The NDC solution was used to make solutions of the following calcium chelators: 1 mM EGTA (Sigma, E4378), 1 mM BAPTA (Sigma, A4926), 1 mM EGTA + BAPTA, and the DMSO carrier control. From these, mucin solutions were made as described above. Adhered synchronized zoospores were pre-treated with the indicated treatment for 10 minutes. The cells were then treated with the matching Bonner’s Salts or mucin solution for 5 minutes before fixation.

### Cell fixation and staining

Cells were fixed with 4% PFA and 50 mM sodium cacodylate, pH = 7.2 on ice for 20 minutes, then for 10 minutes at room temperature before being washed with PEM (100 mM PIPES, pH 6.9; 1 mM EGTA; 0.1 mM MgSO_4_). For fluorescence imaging, cells were stained with 1 μM Tubulin Tracker Deep Red (Thermo Fisher, T34077), 1:1000 calcofluor white (Sigma-Aldrich, 18909), and 5 μM Draq5 (Thermo Fisher, 62251) in PEM buffer for 10 minutes at room temperature. To label actin, cells were permeabilized and stained with 66nM AlexaFluor 488 Phalloidin (Thermo Fisher, A12379) in 0.1% Triton-X 100-PEM buffer for 30 minutes at room temperature.

### Microscopy and Image Analysis

For live imaging, differential interference contrast (DIC) images were captured every 10 seconds using an inverted microscope (Ti-2 Eclipse; Nikon) with a 100X 1.45 NA oil objective, for a period between one minute before and nine minutes after treatment. Timelapse images were analyzed using Fiji (*31*). Cells were considered pre-encysted if the cell had no flagellum and was rounded in the first image. Cells were marked as encysted on subsequent images when the flagellum had completely retracted into the cell body.

Fixed cells were imaged on an inverted microscope (Ti-2 Eclipse; Nikon) with either a 40X 1.45 NA oil objective or a 100X 1.45 NA oil objective and using NIS Elements software. Images were taken using both DIC microscopy and widefield fluorescence microscopy with 360 nm to visualize calcofluor white, 460 nm to visualize phalloidin, and 520 nm to visualize either Draq5 or Tubulin Tracker. IRM images were taken using a SRIC filter cube (Nikon). Images were quantified using automatic thresholding of calcofluor white or Draq5 staining in Nikon Elements, or by hand using the CellCounter Fiji plugin (Kurt De Vos, https://imagej.nih.gov/ij/plugins/cell-counter.html). Unpaired t-tests and ordinary one-way ANOVAs were used to assess statistical significance.

Image processing and analysis for the effect of actin inhibition on *Bd* actin structures (**Fig S3B**) experiments was performed using Fiji, and blind scoring using the CellCounter Fiji plugin. *Bd* zoospores were categorized based on the actin structures present in each cell, as previously defined (*7*): pseudopods (actin staining 1-2 μm wide), actin spikes (actin staining <1 μm wide and ≥1μm long), cortical actin (actin staining along the edge of at least 50% of the cell), actin patches (≥10 actin spots, <1 μm in diameter), or combinations of these. Non-mucin treated cells were normalized to the DMSO-Bonner’s control treatment, and mucin treated cells were normalized to the DMSO-Mucin control treatment due to the effect of DMSO on some actin structures (*7*). One-way ANOVAs were performed on these normalized values and their respective control, for three independent experiments.

Line scan analysis was performed in Fiji using the freehand selection tool and the multiplot function. The outline of control and CytoD treated cells in the presence or absence of mucin were drawn in the actin channel using a line width of three pixels. Line scans along this line were taken in both the actin and the cell wall channel, thus having two identical lines measuring the intensities from two different fluorescent signals. Each line was normalized as a percentage of the highest intensity value. Correlation analysis between the intensity values of actin and the cell wall for each cell was calculated using Pearson’s Correlation Coefficient and plotted with ViolinSuperPlots (*29*), and compared using unpaired t-tests.

Analysis on the relationship between cell wall presence and flagellar retraction was done using Fiji and the Cell Counter Plugin. Ethanol, LatB, DMSO and Jasp treated cells in the presence or absence of mucin were categorized by presence or absence of cell wall and one of four flagellar positions: in, out, partially retracted, not present. Statistical significance was assessed using unpaired t-tests.

## SUPPLEMENTARY DATA

**Data S1. Number of cells analyzed for each biological replicate of each experiment.** The table lists the corresponding figure panel (1st column), treatment conditions (2nd column), replicate number (3rd column), the number of cells analyzed (N; 4th column), and the time point from which the data was collected (5th column). Non-time course experiments have “0” in column 5.

**Movie S1**. DIC microscopy of *Bd* zoospores adhered to ConA for 1 minute before exposure to buffer (+Mock; left) or amphibian mucus (+Mucus; right) for 9 minutes. Images were taken every 10 seconds; movie at 7 frames per second. Time is displayed in minutes:seconds.

## ACKNOWLEDGMENTS

We thank Fritz-Laylin lab members and Sam Lord, Madelaine Bartlett, and Meg Titus for comments on the manuscript, and Edgar Medina for *Spizellomyces* zoospores used in Fig. S2C.

## Author contributions

Conceptualization: L.K.F.-L., K. A.R.; Methodology: L.K.F.-L., K.A.R., E.H.C.G; Formal Analysis: L.K.F.-L., K.A.R, S.M.P.; Investigation: K.A.R, S.M.P.; Writing – Original Draft: L.K.F.-L., K.A.R; Writing – Review and Editing: L.K.F.-L., K.A.R, S.M.P, E.H.C.G.; Visualization: L.K.F.-L., K.A.R, S.M.P; Supervision: L. K.F.-L.; Project administration: L.K.F.-L.; Funding acquisition: L.K.F.-L.

## Data and materials availability

All data are available in the manuscript and the supplementary materials.

---

**Figure S1.**
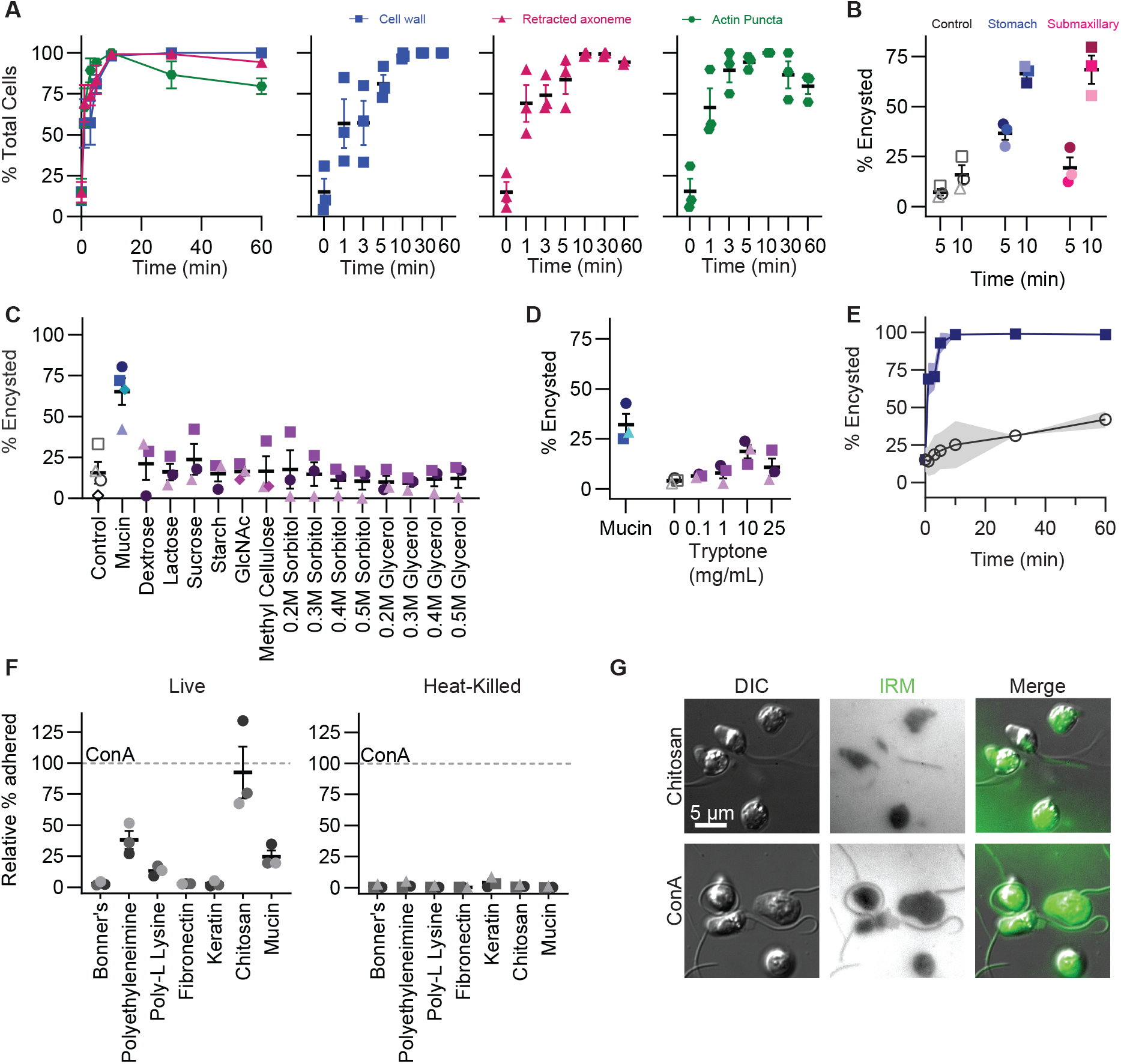
Adherent *Bd* zoospores encyst upon exposure to mucin. (**A**) The quantification of the characteristics of mucin-induced encystation over time after addition of mucin with mean and error shown: axoneme retraction (pink triangles), actin puncta (green circles), presence of a cell wall (blue squares). Biological replicates with mean and standard error are shown below the top graph. (**B, C, D**) The percentage of cells encysted after 5 minutes of exposure to the indicated treatment. The shape corresponds to matching control biological replicate. (**E**) Time course showing the percent of cells with cell walls after treatment with buffer (gray) or 100 mg/mL mucin (blue) over time. The shading shows standard error. (**F**) The percent of cells that adhere to the indicated surface coating, normalized to ConA. Left shows live cells, and right are heat-killed cells. (**G**) Interference reflection microscopy (IRM) showing regions of Bd zoospores in close proximity to a coverslip coated with ConA (top) and chitosan (bottom). (B-E) Open shapes represent the control and filled shapes mucin (blue or pink) or alternative inducers (purple). (A-F) The mean of three independent biological replicates (shapes) is represented by thick black lines, and the standard error by error bars.

**Figure S2.**
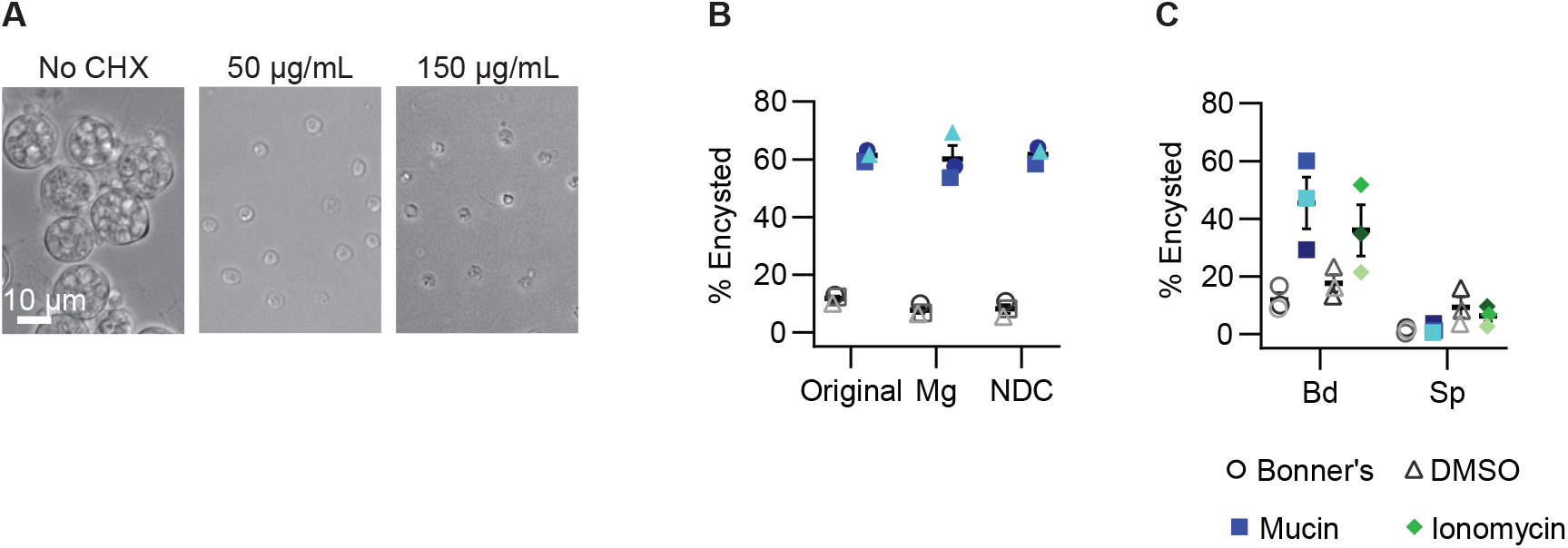
Calcium can induce encystation of *Bd* but not *Sp*. (**A**) Representative DIC microscopy of *Bd* zoospores grown for 72 hours in 1% tryptone media supplemented with cycloheximide (CHX). Note that control cells have developed into mature sporangia (no CHX, left), while cells treated with CHX have ceased development after encystation (middle, right) (**B**) The percentage of cells in original Bonner’s Salts, modified Bonner’s made with MgCl2 instead of CaCl2, and Bonner’s lacking both divalent cations (NDC) that encyst upon exposure to buffer alone (open shapes; gray) or 10 mg/mL mucin resuspended the same buffer (filled shapes; blue). This is a control experiment for the calcium chelator experiments shown in Fig. 2C. (**C**) The percentage of cells that encyst upon exposure to control buffer (circles), mucin (squares), DMSO (triangles), or ionomycin (diamonds) in *Bd* and *Spizellomyces punctatus*, *Sp*. (B-C) Three biological replicates with mean and standard error.

**Figure S3.**
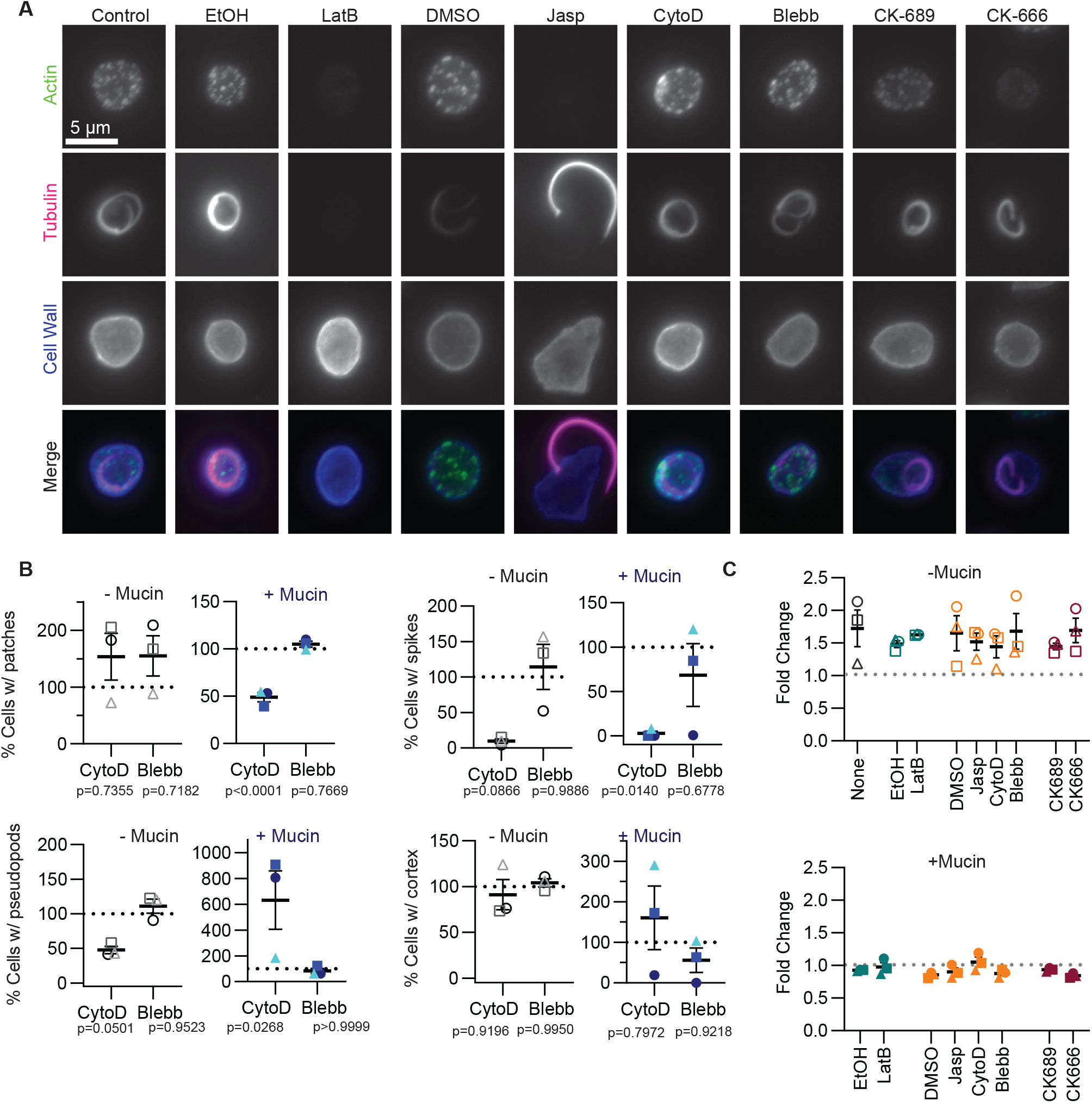
Disrupting actin dynamics alters mucin-induced encystation. (**A**) Representative fluorescence microscopy images of mucin-treated cells exposed to actin inhibitors or matched vehicle controls, stained for actin (phalloidin), tubulin (tubulin tracker), and cell wall (calcofluor white). Image LUTs matched to the “control” cell. (**B**) Percentage of cells with actin structures (patches, spikes, pseudopods, cortex) normalized to DMSO control after treatment with Cytocholasin D or Blebbistatin. Quantification of each actin structure is separated by exposure to buffer (gray circles; left) or 10 mg/mL mucin (blue squares; right). Three biological replicates with mean and standard error. Significance determined by one-way ANOVA. (**C**) The fold change of cell wall intensity, normalized to the median cell wall intensity of the cells labeled “None” exposed to mucin, upon exposure to buffer (top) or mucin (bottom) after treatment with actin inhibitors or matched carrier controls (listed first). (B-C) Replicates with mean and standard error are indicated by circle, square, triangle; open shapes are exposed to buffer and filled to mucin.

